# Stability of Human Serum Amyloid A Fibrils

**DOI:** 10.1101/2020.09.10.291948

**Authors:** Wenhua Wang, Ulrich H. E. Hansmann

**Affiliations:** Department of Chemistry & Biochemistry, University of Oklahoma, Norman, Oklahoma 73019, United States

**Keywords:** Serum amyloid A, Fibril stability, Molecular Dynamics

## Abstract

In systemic amyloidosis, Serum amyloid A (SAA) fibril deposits cause widespread damages to tissues and organs that eventually may lead to death. A therapeutically intervention therefore has either to dissolve these fibrils or inhibit their formation. However, only recently has the human SAA fibril structure be resolved at a resolution that is sufficient for development of drug candidates. Here, we use molecular dynamic simulations to probe the factors that modulate the stability of this fibril model. Our simulations suggest that fibril formation starts with the stacking of two misfolded monomers into metastable dimers, with the stacking depending on the N-terminal amyloidogenic regions of different chains forming anchors. The resulting dimers pack in a second step into a two-fold two-layer tetramer that is stable enough to nucleate fibril formation. The stability of the initial dimers is enhanced under acidic conditions by a strong salt bridge and side-chain hydrogen bond network in the C-terminal cavity (residues 23 - 51) but not affected by the presence of the disordered C-terminal tail.

**Table of Content Graphics:** 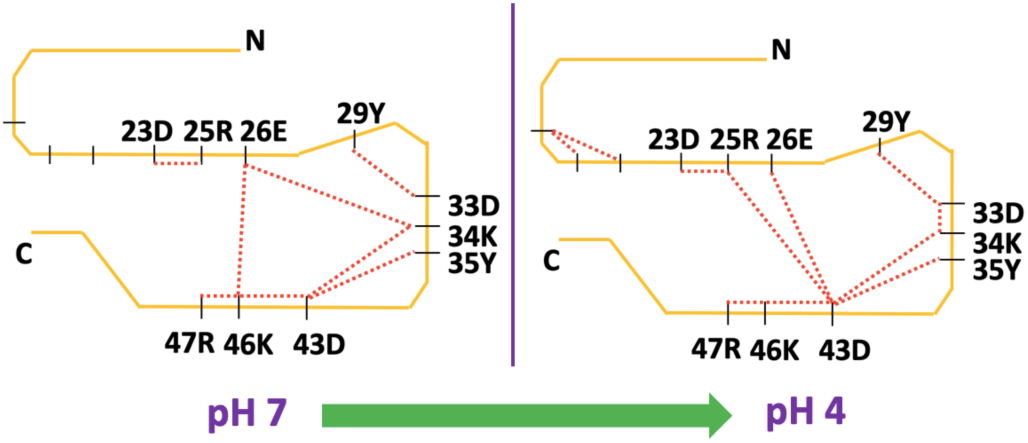

## Introduction

Many diseases are correlated with the presence of ordered fibril-like structures, so called amyloids, that are formed by mis-folded proteins. For instance, in the case of Alzheimer’s disease are these amyloids, here made of Aβ peptides and tau-proteins, extensively studied ^1-4^ and found in the brain. However, amyloid deposits are not always localized. For example, in Serum Amyloid A (AA) amyloidosis is the damage to tissues and organs spread all over the human body.^5, 6^ This *systemic amyloidosis*, which besides humans also appears in more than 50 other species,^7, 8^ is connected with overexpression of serum amyloid A (SAA), as sometimes occurring in inflammatory bowel disease, rheumatoid arthritis, or certain cancers. ^9, 10^ As a consequence of the primary disease can the concentration of acute-phase SAA be 1000 times higher than seen under healthy conditions.^11^ The resulting elevated risk for misfolding and aggregation leads in some cases to AA amyloidosis as a secondary disease that adds to the pathology of the primary disease and complicates its treatment.

In its active form, the 104-residue long SAA assembles into a hexamer and is involved in cholesterol transport and the regulation of inflammation.^12^ On the other hand, SAA fibrils are not formed by the full-sized protein, but are associations of shorter fragments that are derived after enzymatic cleavage. The most commonly found fragment is SAA _1-76_. We have studied in previous work^13^ the conditions that lead to decay of the hexamer, cleavage of the monomers, and their subsequent misfolding. We could propose a mechanism for downregulating SAA activity in case of overexpression that relies on the cleavage of the protein as a way to shift the equilibrium from hexamer to the monomer.^13^ At neutral pH is a structure (coined by us helix-weakened) dominates that can be easily degraded lowering quickly the SAA concentration. However, the N-terminal helix has in helix-weakened configurations a higher risk to unravel, and to release the first eleven residues. These N-terminal residues are known to be amyloidogenic^14-16^ can then form a strand-like segment that in turn may nucleate amyloid growth. If acidic conditions raise this risk, the equilibrium therefore shifts toward the second motif (termed by us helix-broken) where the N-terminal residues stay cached in a helix, making the SAA_1–76_ fragment more difficult to degrade, but also less aggregation-prone.

The above described mechanism assumes a competition between fast degradation of SAA fragments and their tendency to form amyloids. Amyloidogenic conditions are countered by a shift to a more stable and less easily degradable form, often avoiding amyloidosis in cases where the primary disease causes overexpression of SAA. However, the equilibrium between amyloids and free SAA fragments depends not only on the relative frequency of the two fragment motifs but also on the stability of the final fibrils. Hence, possible treatment options could involve either stabilization of the N-terminal segment in the SAA_1-76_ fragments (decreasing the risk to nucleate amyloid formation), or destabilizing the fibrils. The present work is motivated by the second avenue and explores the factors that drive fibril formation and stabilize the fibril geometry.

The kinetics of fibril formation is difficult to probe in either experiments or in silico. Computationally more accessible^17, 18^ is the stability analysis of fibrils which, however, requires a structural model of the fibril. The human SAA fibril model was only recently resolved by electron cryo-microscopy,^19^ and is deposited in the Protein Data Bank (PDB) under identifier *6MST*. Because of experimental difficulties only the fragment SAA_2-69_ (shown in Figure 1) was resolved, and here only residues 2-55 have a defined structure while residues 56-69 are disordered.

**Figure 1.**
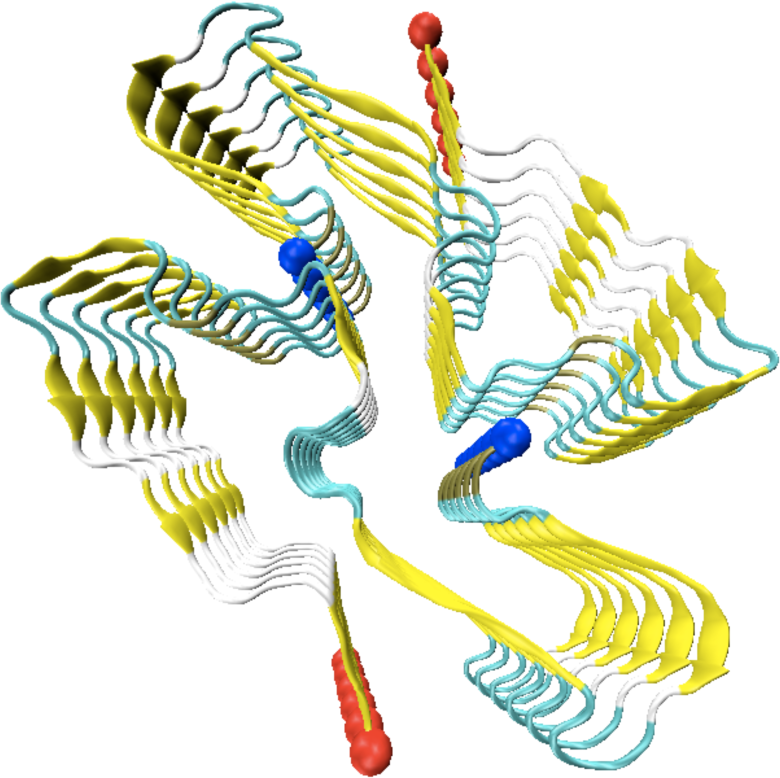
Cryo-EM model of the human SAA fibril (PDB ID: 6MST). The N- and C-termini are marked by blue and red spheres, respectively. The β-strands are colored in yellow, turns in cyan and coil regions in gray.

The ordered part of the human SAA fibril adopts a right-hand twist, with each chain forming seven β-sheets, β1-β7. The N-terminal region is made of about 20 residues and is stabilized by hydrophobic interaction involving six hydrophobic residues (F3, F5, F10, M16, Y17, and A19). Unlike the N-terminal region, the C-terminal region of human SAA fibril is stabilized by the ionic interactions among the charged residues (D23, R25, E26, K34, D43, and R47) buried inside a C-terminal cavity. Two proto-fibrils are packed together, with the packing interface stabilized by ionic, hydrophobic, and polar cross-stack interactions. The center of the interface is a so-called “steric zipper”, formed by the packing of β3-strands (residues Y29 and I30) of chains on opposite proto-fibrils. A salt-bridge connecting the N-terminus of a chain in one layer of a protofibril with residue D33 of a chain of the same layer in the opposite proto-fibril is also stabilizing the packing interface, ^19^ see Figure 1. The tight packing of the protofibrils makes us wonder whether packing interactions between two chains may already lead to a stable nucleus, i.e., whether fibril formation starts with nucleation and elongation of single protofibrils, or whether both protofibrils grow out of such dimers kept together by packing interactions. Hence, the first question that we pose is that for the smallest stable fibril fragment, i.e., the nucleus which starts fibril growth, and whether this fragment is stabilized by stacking or packing interactions. As it is known that acid conditions further SAA fibril formation, we ask as a second question how acidic conditions affect the fibril stability. Finally, in our third line of research we study the role of the first eleven residues (an amyloidogenic region) and of the residues 56-76 (a disordered region) for fibril stability. We study the later region because SAA_1-76_ (and not SAA_1-55_) is the most commonly found fragment in human SAA fibrils. Hence, it appears to be likely that the disordered region, which we assume to extend to residue 76, plays also a role in fibril formation and fibril stability. ^19^

Analyzing our molecular dynamics simulations we conclude that the critical size for fibril stability is a two-fold two-layer tetramer. We conjecture that stacking within the same fold precedes packing between two folds. Acidic conditions are most important at early stages of fibril formation. By altering the salt-bridge network for the C-terminal cavity, they encourage the stacking of chains, initiated by N-terminal regions of different chains forming anchors. As shown in our previous work,^13^ these N-terminal residues are more easily released from the N-terminal helix in the native structure once the C-terminal residues 65-76 become disordered. However, the disordered C-terminal tail does not affect by itself the stability of the SAA fibril geometry.

## Materials and Methods

Models used in our simulations are generated from the cryo-EM structure as deposited in the Protein Data Bank (PDB) under identifier 6MST. The fibril structure is made of SAA_2-55_ fragments, i.e., without the first residue which we also do not consider in our investigation. The model is a two-fold six-layer system, from which we have successively deleted chains in the bottom and top end layer to obtain a sequence of models with two-fold three-layers (F2L3), two-fold two-layers (F2L2), and two-fold one-layer (F2L1). By deleting one of the folds, corresponding single-fold models are generated, i.e., one-fold three-layers (F1L3), one-fold two-layers (F1L2), and one-fold one-layer (F1L1) systems. In order to test the role of the first eleven N-terminal residues we generated also a series of truncated models (F1L1-,F1L2-,F1L3-,F2L1- and F2L2-) where these residues where deleted from the chains. In another series of models (F1L1A, F1L2A, F1L3A, F2L1A, F2L2A, F2L3A) we accounted for the effect of acid conditions in an approximated way by altering the protonation states of residues E9, E26, H37 to the one expected at pH=4 while these protonation states correspond in the parent models to pH=7. Each of the single-fold models is placed in a cubic box of edge size ∼ 9.6 nm and filled with ∼ 28000 water molecules, while for two-fold models the box has an edge size of ∼ 10.4 nm and is filled with ∼ 37000 water molecules. Since most fibrils taken from patients are made from the larger SAA(1-76) fragments, we have in addition generated one model where the residues 56-76 were added. These residues are not resolved in the cryo-EM structure indicating that that this segment is unstructured. We generated this specific model only after we had determined first the two-fold two-layer F2L2 as the smallest stable system. Hence, our extended model F2L2+ is also a two-fold two-layer system, with the added additional residues 56-76 assuming a coil configuration. The extended model is again placed in a cubic box of edge size of ∼ 17.5 nm that is filled with ∼ 74000 water molecules. All simulated models are listed in Table 1.

**Table 1.**
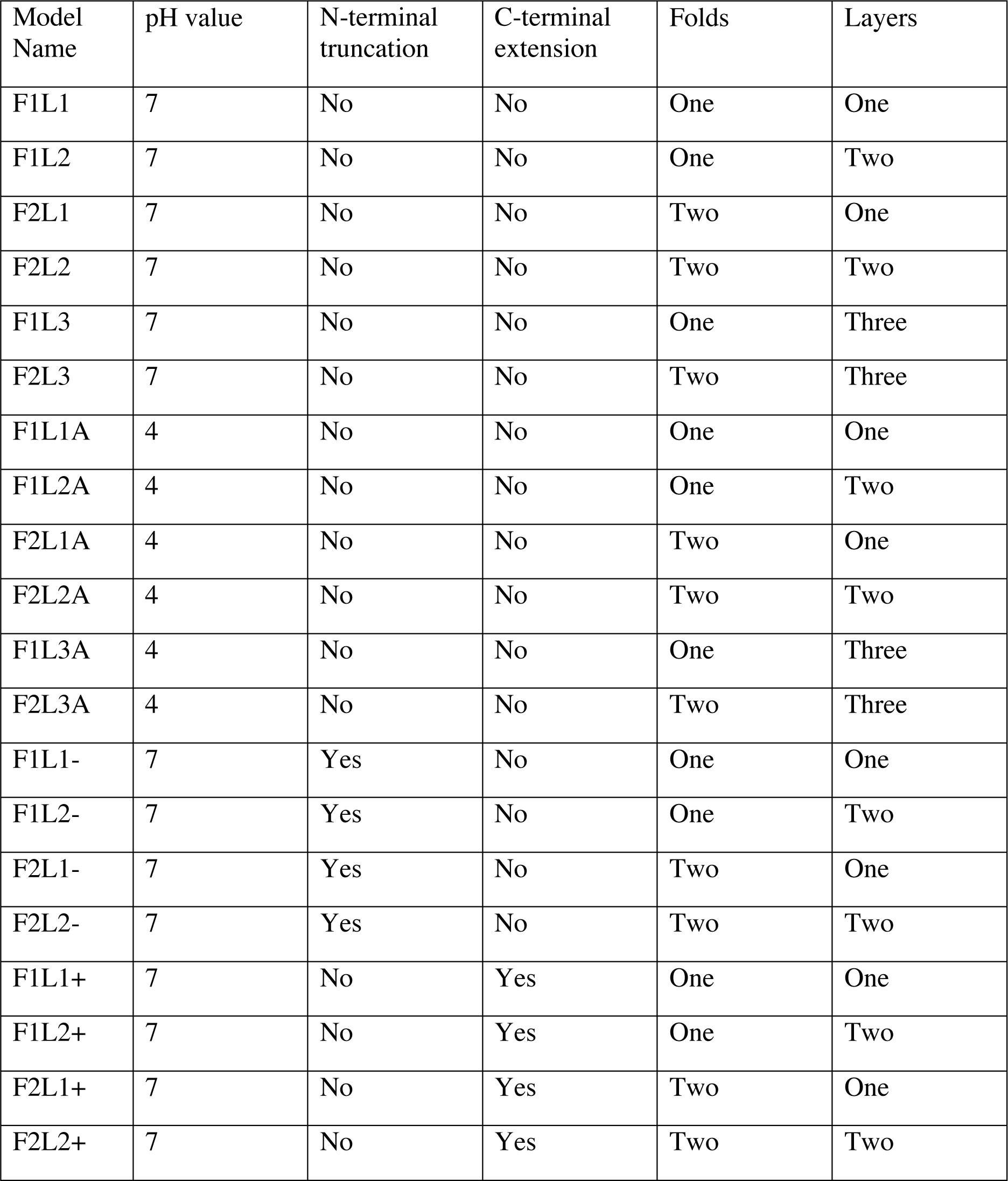
List of the considered models.

Our simulations rely on the GROMACS 2018.2 software package. ^20^ We use the CHARMM36m force field to describe interactions between atoms, and TIP3P for modeling water. ^21, 22^ The LINCS and SETTLE algorithms are used to constrain bond length and water geometry. ^23, 24^ As we utilize periodic boundary conditions, we rely on the Particle mesh Ewald algorithm (with a cut-off of 1.2 nm) to calculate electrostatic interactions. The same cut-off is used in calculations of van der Waals interactions. Molecular dynamics simulations are done in an isothermal-isobaric (NPT) ensemble, ^25^ with the temperature set to 310 K by a v-rescale thermostat and pressure to 1 Bar by a Parrinello−Rahman barostat. ^26, 27^ Three independent trajectories of 100 ns length are generated for each system with the exception of the extended systems which were simulated only for 80 ns (F1L2+ and F2L2+) or 30ns (F1L1+ and L2L1+). Each of the three trajectories starts with an initial configurations derived from the above generated models by small random variations of coordinates and the velocity distribution. Data are collected and stored every 50 ps for later analysis. Only the last 70 ns (and in case of F1L2+ and F2L2 the last 50 ns, and for F1L1+ and F2L1+ the last last 50ns) are used of the trajectories are used for our stability analysis, which relies on GROMACS built-in tools and in-house code for measuring quantities such as RMSD, RMSF, Contact map, etc. ^20^ Binding energies are approximated by the MMGBSA approach ^28^ using Amber 12 package and its MMGBSA tools. ^29, 30^

## Results and Discussions

### Critical size of SAA fibril and potential amyloid formation mechanism

In order to study the stability of the recently resolved SAA fibril model, we start by determining the critical size of the fibril, that is the size above which the fibril is stable over extended periods of time while below this size the fibril will decay on the time scale of our simulations. Representative structures from simulations of various fibril fragments built from SAA_2-55_ – chains are shown in Figure 2. The simulations were done at neutral pH and the configuration taken at the end of one of the three 100 ns long trajectories of fibrils. The evolution of the fibrils is quantified by the plots of the root-mean-square deviation (RMSD) to the start configuration as function of time in Figure 3a. The presented values are calculated separately for each chain, and then averaged over all chains in the fibril. We show only one of the three trajectories, namely the one that let to the largest final RMSD value (i.e. the trajectory leading to the largest change). Corresponding average Root-mean-square-fluctuations (RMSF) of the residues in the chains, a measure for the flexibility of the chain at the point of the specific residue, are shown Figure 3b.

**Figure 2.**
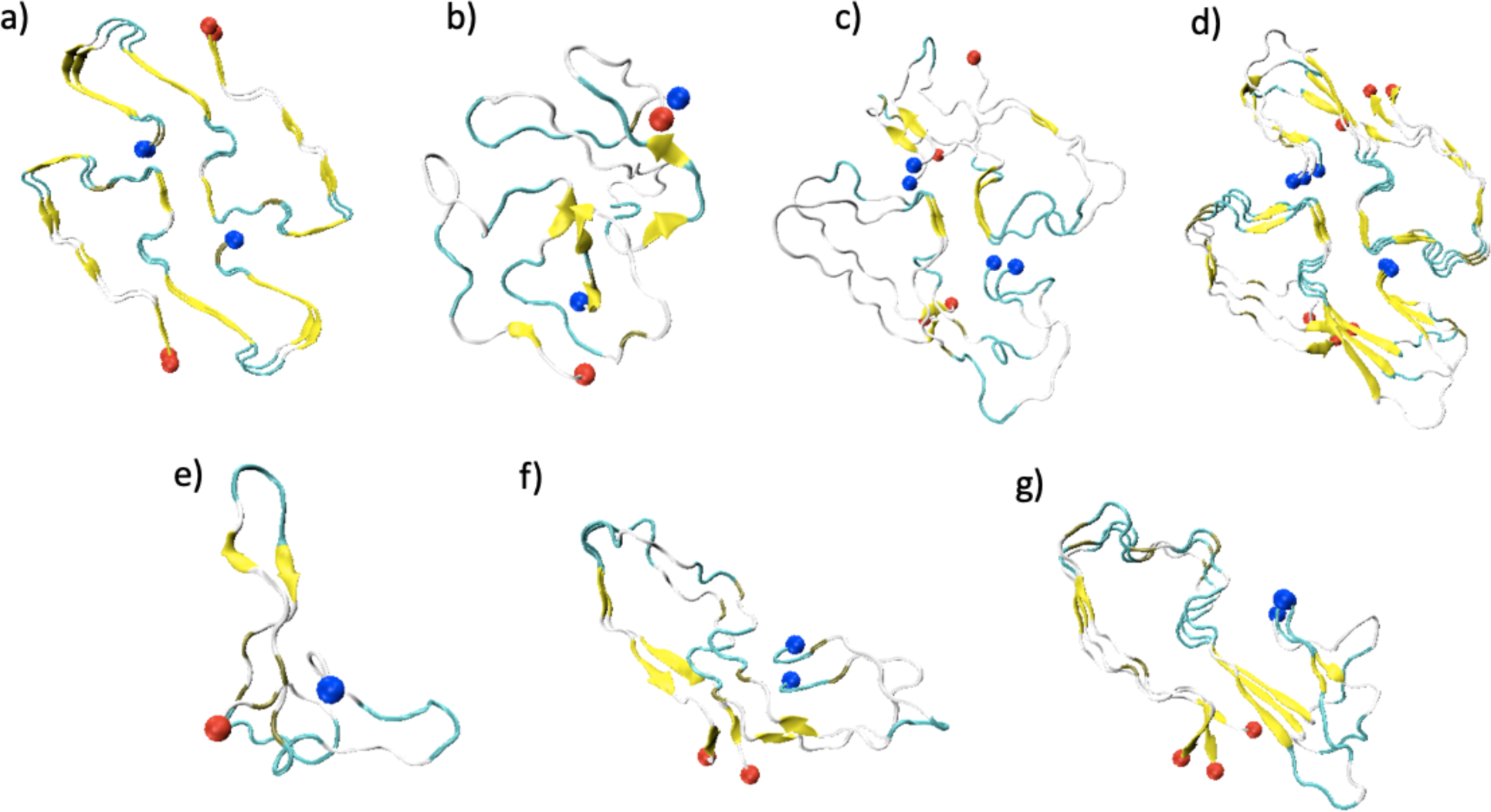
a) Initial structure of the fibril fragment F2L2. All simulations of fibril models built from SAA_2-55_ chains start from configurations were the individual chains have the same structure as in this model. Representative configurations, taken at the end of one of the three 100 ns long trajectories of fibrils simulated at pH=7, are shown in b) -g). Specifically, final configurations are shown for F2L1 (b), F2L2 (c), F2L3 (d), F1L1 (e), F1L2 (f) and F1L3 (g). *N- and C-*terminus are marked by blue and red spheres, respectively. The β-strands are colored in yellow, turns in cyan and coil regions in gray.

**Figure 3.**
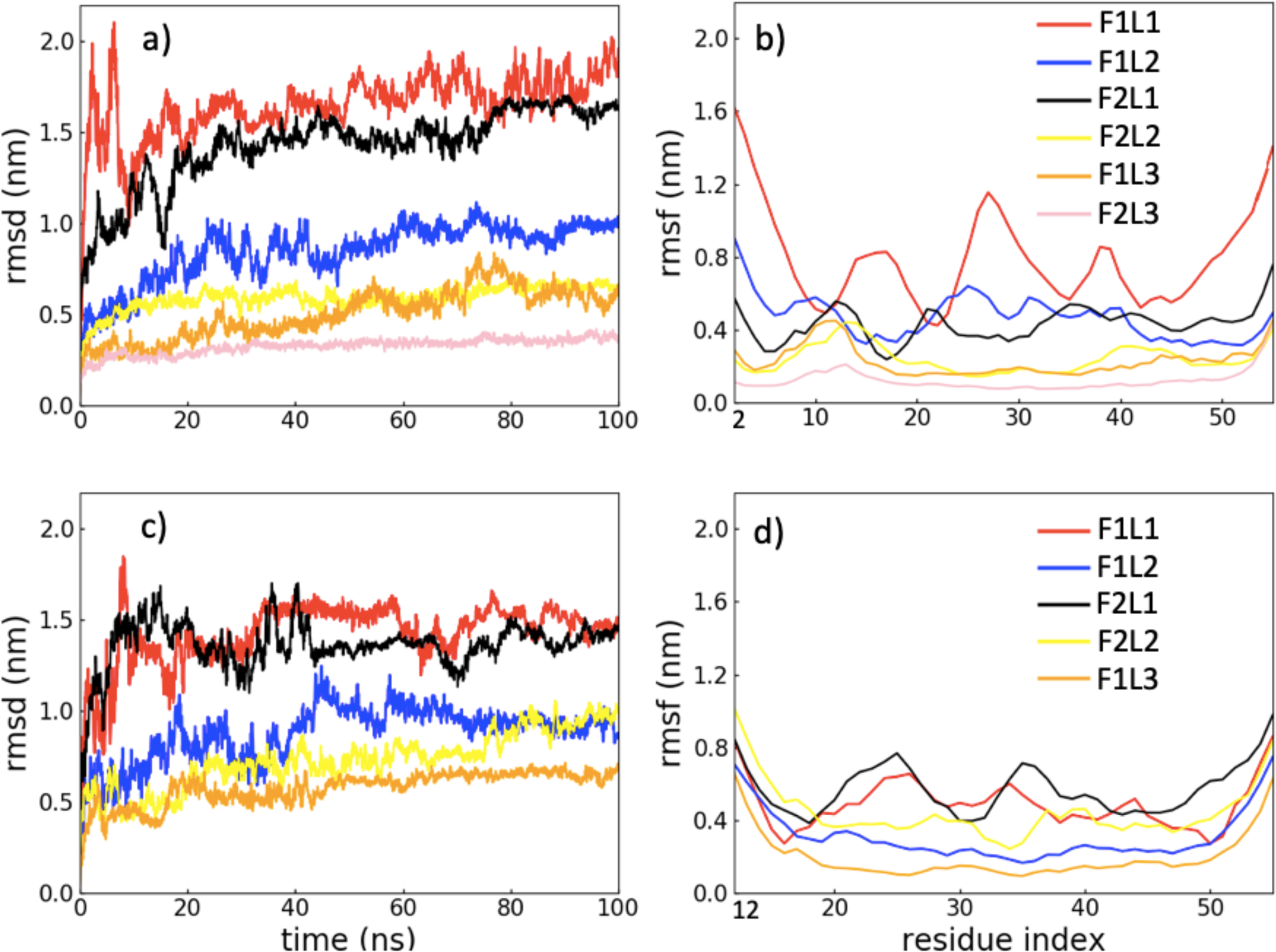
a). Average RMSD per chain for various fibril models simulated at pH=7. Only the trajectory leading to the highest final RMSD value is shown. The trajectories are for the F1L1 (red), F1L2 (blue), F2L1 (black), F2L2 (yellow), F1L3 (orange) and F2L3 (pink). Corresponding RMSF values are shown in b). In c) and d) we show the same quantities measured in simulations of the corresponding truncated systems: F1L1-(red), F1L2-(blue), F2L1-(black), F2L2-(yellow), F1L3-(orange), F2L3-(pink).

Comparing the various models, we find that at neutral pH the SAA_2-55_ monomer F1L1 and the two-fold one-layer dimer F2L1 are unstable. This observation is confirmed by visual inspection of the final configurations in Figure 2b and 2e. Hence, our data exclude not only SAA_2-55_ monomers as possible nucleus for fibril formation, but also dimers made of two chains packed together in one layer. On the other hand, the stability of the models increases with the stacking of layers. The same pattern is seen in table 2 where we show the change in solvent accessible surface area (SASA) over the length of the trajectory, with large changes for the monomer (F1L1) and the F2L1 dimer, and very little changes for the F2L2 and larger systems. We remark that the SASA values are divided by the number of residues in order to compare systems with different number of chains.

**Table 2.** Change in solvent accessible surface area (SASA) as measured in all trajectories obtained from simulations of SAA fibril fragments at neutral pH. Compared with the start configuration are averages taken over the last 20 ns. The shown values are obtained by dividing by the total number of residues. This allows us to compare systems of different number of chain and chain sizes.

|  | F1L1 | F1L2 | F2L1 | F2L2 | F1L3 | F2L3 |
| --- | --- | --- | --- | --- | --- | --- |
| | $\Delta\text{SASA} (\text{\AA}^2)$ | | | | | |
| Run-1 | -34.5 | 6.2 | -41 | 2.7 | 7.3 | 1.3 |
| Run-2 | -19.6 | 8.8 | -42.2 | 2.5 | 7.2 | 2.6 |
| Run-3 | -31.1 | -2 | -30.9 | 0.8 | 3.9 | 4.9 |
| Average | -28.4 (9.5) | 4.3 (5.3) | -38 (5.7) | 1.9 (1.6) | 6.1 (2.1) | 2.9 (1.8) |

Note that the one-fold two-layer model F1L2, i.e., a dimer made of two chains stacked on each other, has higher stability than the two-fold one-layer dimer F2L1, where the two chains are in packed together in the same plane. This can be seen from both visual inspection, see the representative configurations in Figure 2f, and from the change in solvent accessible surface area (SASA) over the length of the trajectory, as shown in table 2. While the change for F2L1 is even larger than for the monomer (F1L1), it is much smaller for the F1L2 dimer where the two chains are stacked on top of each other. The variation of the RMSD values measured for the three trajectories, shown in Figure 4a (F1L2) and 4b (F2L2), and visual inspection, see the representative structure in fig 2f, indicate that the F1L2 dimer is still not stable enough to serve as nucleus for fibril growth. However, it may serve as a metastable state on the way to forming the two-fold two layers system F2L2, the smallest assembly seen in our simulation that is stable enough to nucleate fibril elongation.

**Figure 4.**
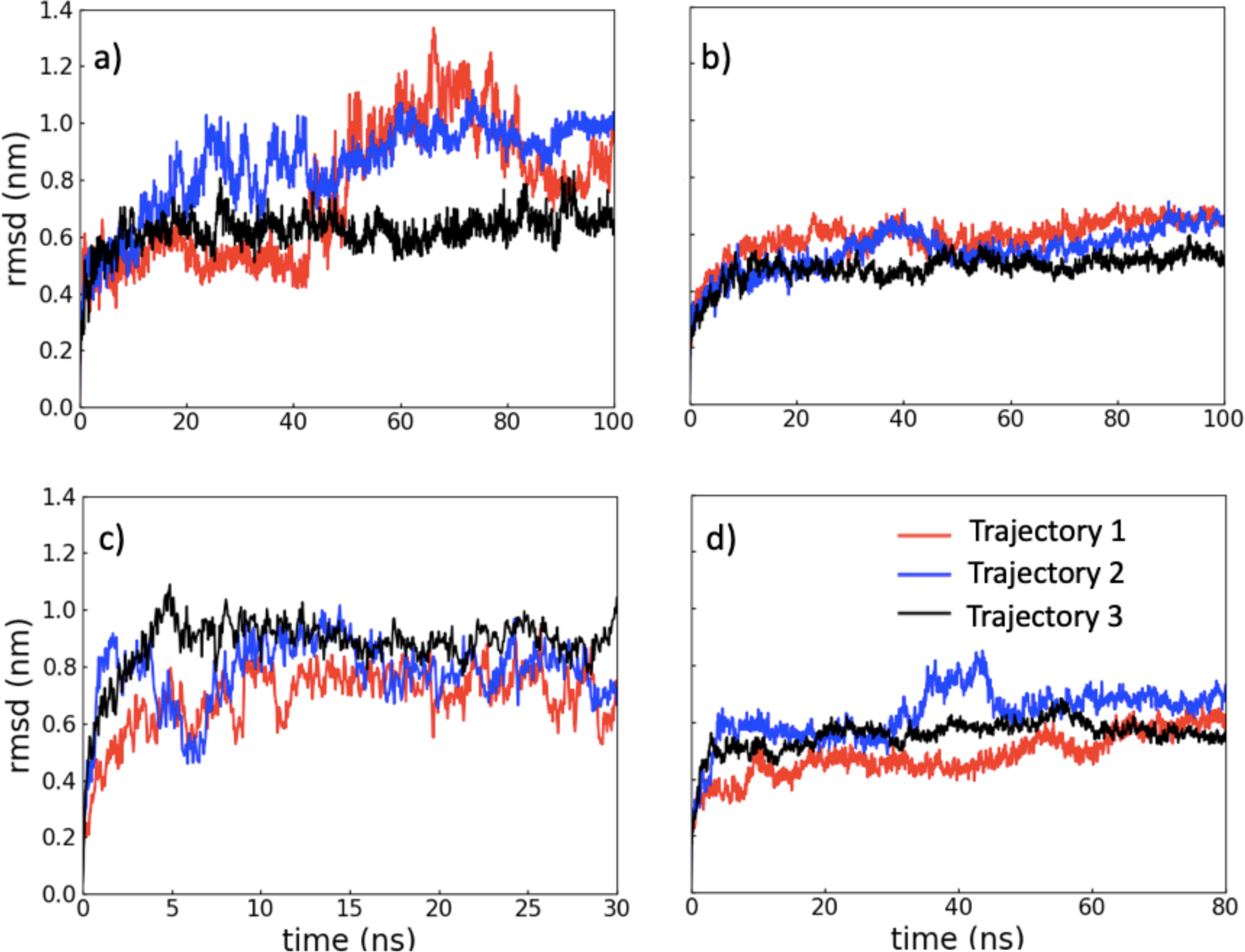
Average RMSD per chain values for each of the three trajectories simulated for the SAA_2-55_ fibril models a) F1L2, b) F2L2, and for the elongated SAA_2-76_ models F1L2+ (c) and F2L2+ (d).

Hence, our stability analysis of the various models suggests that at neutral pH the formation of the critical nucleus starts with two monomers forming mainchain hydrogen bonds and stacking into a metastable dimer, with each chain assuming a fibril-like configuration. In the PDB fibril model, such stacking reduces the solvent accessible surface area by approximately 48 Å^2^ per residue at the interface between two chains. Packing of two such dimers with their N-terminal region leads to a tetramer where the chain configurations are sufficiently stabilized by the interactions between the chains and an additional reduction of about 12 Å^2^ per residue. However, more indicating for the gain in stabilizing hydrophobic forces is a comparison of the SASA values of the thermalized structures as averaged over the last 20 ns of each trajectory for the respective systems, shown by us in table 3. Stacking of two chains leads to an effective reduction of SASA of 15.2 (7.7) Å^2^ per residue for F1L2, and packing of two F1L2 dimers further reduces it by another 10 (4) Å^2^ per residue. However, not only hydrophobic forces contribute to the stabilization of fibril fragment. This can be seen by looking into the binding energies as approximated by MMGBSA calculations. For stacking of chains we find a values of -59.4 (22) kcal/mol, while binding for the packing of two F1L2 dimers gives a value of -55 (33) kcal/mol, larger than what is expected from the change in SASA.

**Table 3.** Solvent accessible surface SASA per residue (in Å^2^), averaged over the last 20 ns of all three trajectories of each system.

| Systems | Total | Hydrophobic | Hydrophilic |
| --- | --- | --- | --- |
| F1L1 | 96.1 (9.5) | 51.1 (5.5) | 45 (4.5) |
| F1L2 | 80.9 (5.3) | 42.1 (3.1) | 38.8 (2.8) |
| F1L3 | 65.9 (2.1) | 33.3 (0.9) | 32.6 (1.8) |
| F2L1 | 80.4 (5.7) | 41.9 (4.2) | 38.5 (2) |
| F2L2 | 70.9 (2.6) | 36.5 (2.0) | 34.4 (1) |
| F2L3 | 56.1 (1.8) | 27.6 (0.9) | 28.5 (1) |
| F1L1A | 100.7 (10) | 53.4 (6.1) | 47.3 (4.6) |
| F1L2A | 76.2 (2.6) | 40.4 (1.1) | 35.8 (2.3) |
| F1L3A | 63.5 (1.4) | 32.5 (0.9) | 31 (1.1) |
| F2L1A | 86.9 (8.6) | 47.2 (5.6) | 39.7 (3.4) |
| F2L2A | 67.7 (3) | 34.4 (1.2) | 33.3 (2) |
| F2L3A | 57.6 (1) | 28 (0.8) | 29.6 (0.8) |
| F1L1- | 81.2 (4) | 41 (2.8) | 40.2 (2.4) |
| F1L2- | 67.2 (3.0) | 34.4 (2.4) | 32.8 (1.3) |
| F2L1- | 68.7 (3.4) | 34.2 (1.7) | 34.5 (2.3) |
| F2L2- | 65.2 (2.1) | 32.6 (1.1) | 32.6 (1.3) |
| F1L1+ | 104.5 (10.1) | 56.9 (6.4) | 47.6 (4.3) |
| F1L2+ | 77.6 (2.0) | 40.8 (1.5) | 36.8 (1.8) |
| F2L1+ | 88.8 (5.9) | 46.5 (4.1) | 42.3 (2.2) |
| F2L2+ | 71.3 (1.6) | 35.7 (1.4) | 35.6 (1.4) |

The so derived two-fold two-layers tetramer F2L2 then serves as a nucleus for further fibril growth, see Figure 2 c. With growing number of layers decrease the RMSD and RMSF values of the various systems studied in our simulations. The corresponding values are shown in Figure 3a and 3b. The differences between the observed stabilities are small, and systems larger than the critical size preserve their fibril structure on the time scale of our simulations, i.e., can be considered to be stable, see also the representative final configurations in Figure 2d and 2g. The critical size determined by our simulation is also consistent with previous measurements of the Gibbs free energy needed for dissociation of two protofibrils. Using PDBePISA it was found that a protofibril has to have at least two layers in order for this dissociation energy to be positive.^19^

### Acidic conditions enhance early stage aggregation

The situation is different at pH=4,. We have simulated such acidic conditions in a simplistic way by protonating the sidechains of Glutamic acid and Histidine. While the fibril stability changes similarly with number of folds and layers as in the simulations done at neutral pH, we observe differences. For instance, the root-mean-square-fluctuations (RMSF) of the residues in Figure 5a is smaller for the one-fold two layers system F1L2A than the one for the corresponding system at pH=7, namely F1L2. Differences are most prominent for residues 12 – 37 and for the C-terminal cavity formed by residues 45 – 55. The lower root-mean-square fluctuations imply a higher stability of this assembly than seen under neutral condition. These differences in RMSF and stability are not seen when comparing the two-fold two-layer F2L2A tetramer with the corresponding system at neutral pH, F2L2, see Figure 5b. Hence, while acidic conditions do not raise the stability of fully-formed fibrils, see Figure 5c and 5d, we conjecture that they enhance fibril formation by making it easier to form a nucleus. This is because at pH=4 leads the stacking of two chains to a dimer that is more stable than it is at pH=7. The stronger binding by stacking is related to a larger loss of effective solvent accessible surface are at pH=7. Taking the values shown in table 3, we find a value of -24.5 (7.2) Å^2^ at pH=4 compared to 15.2 (7.7) at neutral pH. While for neutral pH the binding energy (as approximated from MMGBSA calculations) of two monomers to the stacked F1L2 dimer is -59.4 (22) kcal/mol, it is -103.4 (22) kcal/mol at pH=4. The resulting longer lifetime makes it more likely for a F1L2 dimer to find a partner and assemble to the F2L2 tetramer that is the critical nucleus. Note that we do not find the larger binding energies for the F2L1 dimer where the two chains are packed together instead of stacked on top of each other. Here, the corresponding binding energies are -40.7 (11) kcal/mol at pH=7 and -44 (242.2) kcal/mol at pH=4.

**Figure 5.**
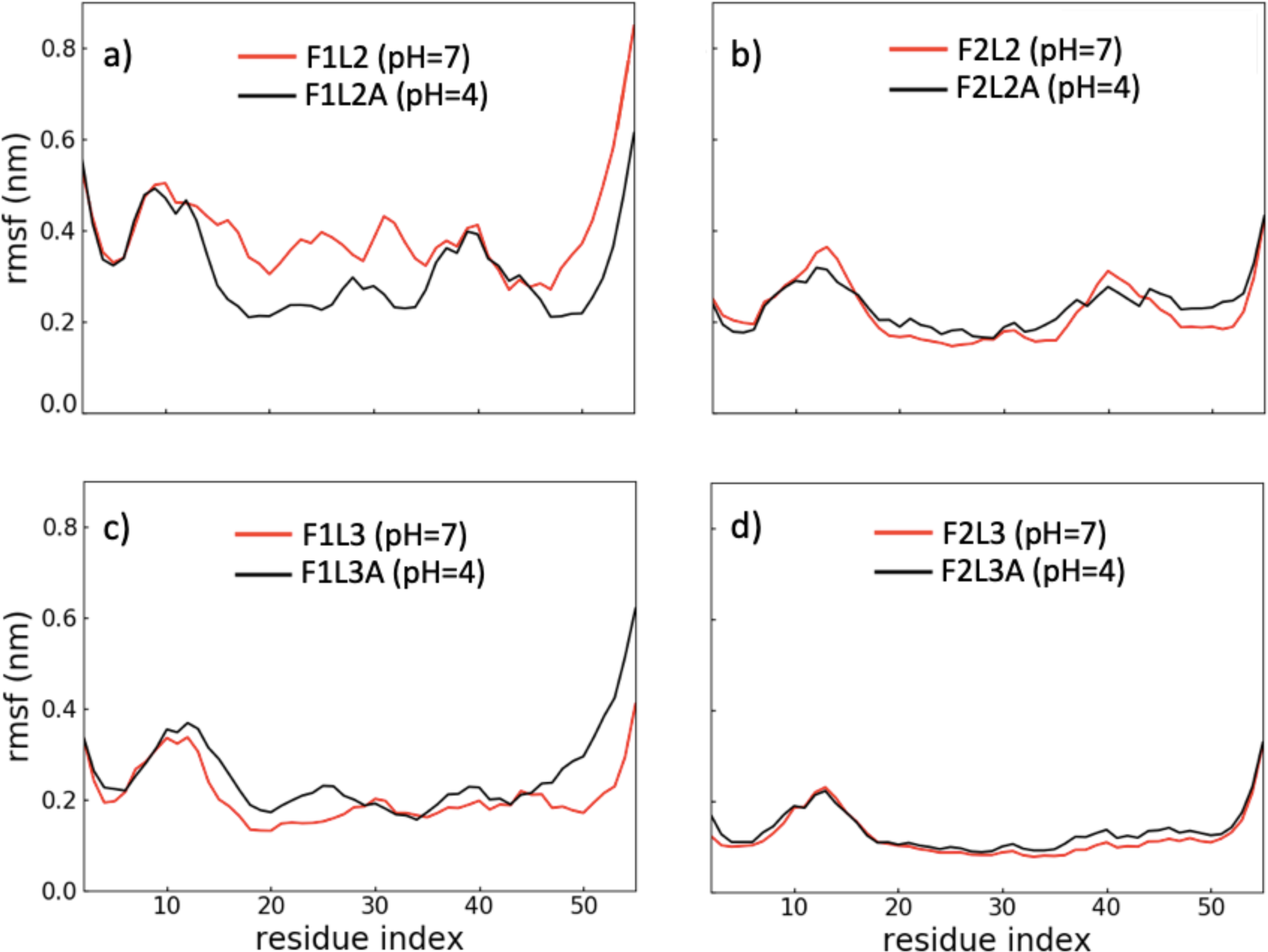
Average RMSF of residues as measured in simulations of fibril models at either neutral pH (black) or under acid conditions (pH=4, drawn in red). In a) we consider the single-fold two-layer F1L2 (F1L2a), in b) the two-fold two-layer F2L2 (F2L2a), in c) the single-fold triple layer F1L3 (F1L3a), and in d) the two-fold-triple-layer F2L3 (F2L3a).

What causes the additional stabilization of the F1L2 dimer under acid conditions? In order to answer this question we have studied how the number and distribution of contacts changes when lowering the pH. We show in Figure 6a the contact map for the single-fold double-layer dimer F1L2 at neutral pH, and in Figure 6b the corresponding map for the same dimer at pH=4 (F1L2A). While at neutral pH only a few long-range side-chain contacts (between F3 and M24) contribute to stability, under acidic conditions are additional contacts found in the C-terminal cavity region. Example are the contacts between residue E26 and the group of residues K46, R47 and G48; and the contacts between the group of residues R25, E26 and K34 with D43. Note especially the role of the glutamic acid E26 which becomes protonated at pH=4. This allows the residue to form now multiple sidechain hydrogen bonds and salt-bridges in the C-terminal cavity (residues 23 - 51) that further stabilize the meta-stable dimer. For instance, at neutral pH salt-bridge interactions between E26 and K34 (8 % occupancy) or E26 and K46 (7 % occupancy) connect the glutamic acid weakly with the two groups of residues 47-K46-D43 and residues Y35-K34-D33-Y29, see Figure 6g). However, at pH=4 interacts E26 preferentially with D43 enabling now also an side-chain hydrogen bond interaction between R25 and D43 (8 % occupancy), and a salt-bridge interaction between K34 and D33 (7 % occupancy). As R25 forms also a contact with D23 (with occupancy of 50 %), and D43 with R47 (with 27 % occupancy), the two groups of residues D23-R25-E26 and R47-K46-D43 become connected, and form now together with the residue group Y35-K34-D33-Y29 a network of intra-chain side chain hydrogen bonds and salt bridges, see Figure 6h. The stronger intrachain contact network in the C-terminal cavity change with lowering the pH, increasing the structural stability of the chain folds.

**Figure 6.**
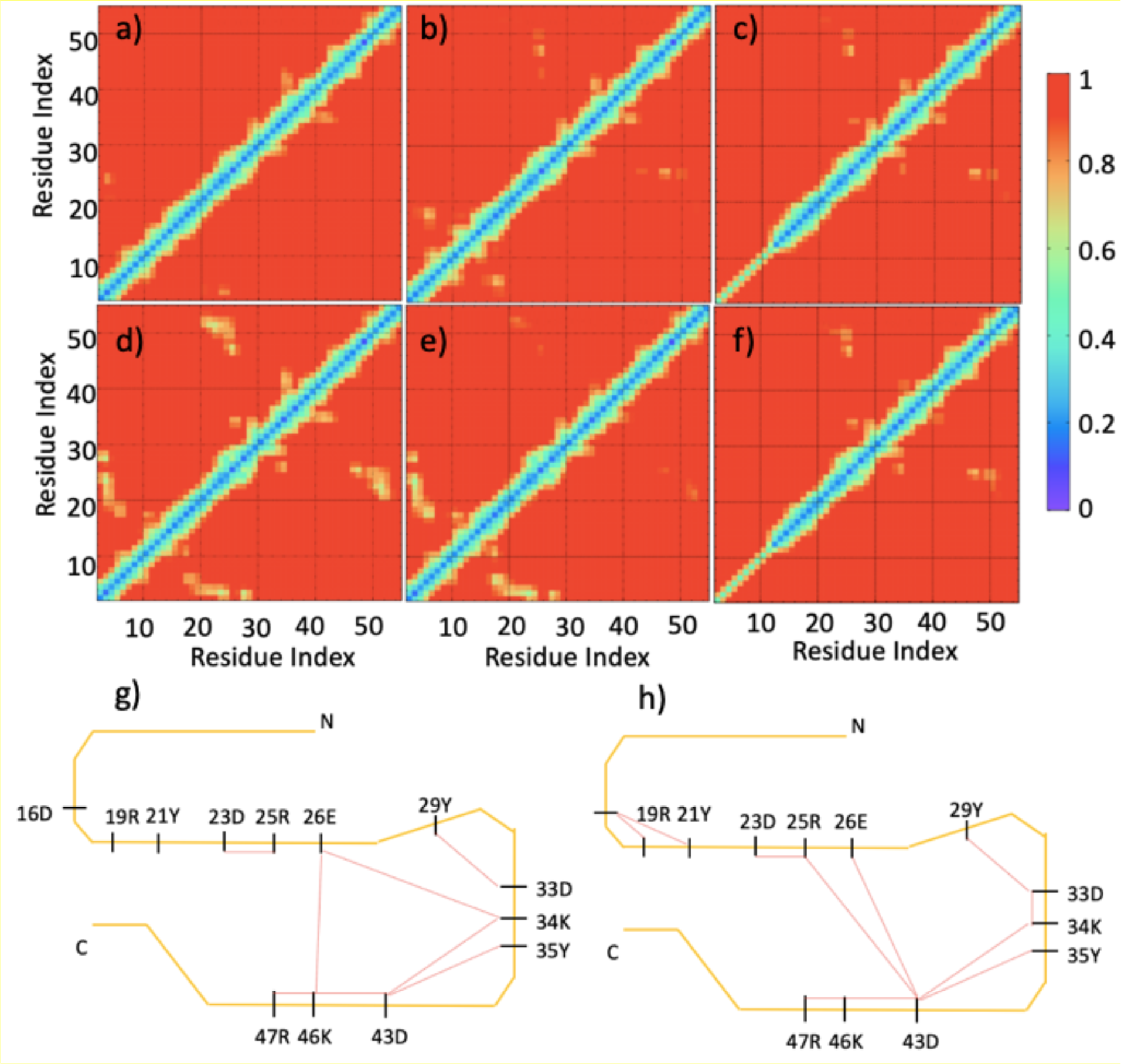
Intra-chain side-chain contact map for the one-fold two-layer fibrils a) F1L2 (pH=7), b) F1L2A (pH=4), c). F1L2-(without N-terminus); and the corresponding two-fold two-layers fibrils d) F2L2, e) F2L2A and f) F2L2-. Contact distance is encoded by coloring. A schematic representation of the contact network is shown for pH = 7 in g) and for acidic conditions (pH=4) in h). Only side chain interactions with an average occupancy per chain larger than 4 % are shown.

On the other hand, the pH-dependence of fibril stability disappears once the critical nucleus size is reached or exceeded. This can be seen, for instance, in Figure 6d and 6e, where the contact map for two-fold two-layer tetramer F2L2 (pH=7) and F2L2A (pH=4) are shown. At pH=7 exist in the tetramer already a multitude of contacts within both the N-terminal and C-terminal regions of SAA chains, and as a consequence, there is little difference in the overall stability when going from neutral to acidic conditions. This picture is similar for larger assemblies, which indicates that once a critical size is reached the already at pH=7 existing intra-chain and inter-chain contacts suffice (unlike for the meta-stable F1L2 dimer) to maintain the stability of the fibril. Hence, acidic conditions do not change the stability of fibrils but encourage their formation by stabilizing the early-stage on-pathway meta-stable F2L2 dimers.

### Roles of N-terminal amyloidogenic region and C-terminal disordered region

The above discussion has already pointed out the importance of inter- and intra-molecular interactions involving the N-terminal residues for maintaining SAA fibril stability. Our observation is consistent with previous experimental studies, which showed that the truncation of the first eleven residues prevents SAA fibril formation. Hence, in order to understand the role of these residues in more detail, we have also considered truncated fragments where the N-terminal residues are missing, and compared their stability with that of the parent fragments. These simulations were done at neutral pH, and RMSD and RMSF values of the truncated system are shown in Figure 3c-3d. The RMSD values as function of time evolve in a similar pattern for the truncated system as for the non-truncated ones for the two-fold systems, but in opposite way for one-fold systems. For example, the RMSD value for F1L1-decreases to ∼ 0.16 nm in the last 20 ns simulation interval compared to F1L1, see fig 3a and 3c. In a similar way decrease the RMSF values in the one-fold systems for the truncated system F1L1-, F1L2-, and F1L3-with respect to non-truncated systems F1L1, F1L2, and F1L3. Again the opposite behavior is observed for two-fold systems in fig 3b and 3d.

For the monomer F1L1- and the single-fold dimer F1L2-leads the lack of the N-terminal region to a more rigid C-terminal region, see Figure 7a and 7b. This rigid C-terminal region is stabilized by the side-chain interactions (fig 6c) and its rigidity implies a higher stability of these fibril models. This is also seen in a binding energy (as approximated by MMGBSA calculation) that with -70.8 (20.5) kcal/mol has a larger magnitude than the non-truncated system (59.0 (20.9) kcal/mol). These results appear to contradict experimental observations that SAA cannot form fibrils when the first eleven residues are truncated. However, our results can be reconciled with the experiments if one assumes that the SAA fibril form by a process where the N-terminal regions serve as anchors, that ease the stacking of the rest of chains. Once the stacking of two chains is completed, they will be stabilized not only by interactions between N-terminal residues, but in addition also by such involving the C-terminal residues, see Figure 6c and 6f. Our hypothesis implies that existing fibrils will not dissolve if the first eleven residues are cleaved, but that fragments derived from such fibrils with truncated chains cannot nucleate fibril growths and elongation. While going beyond the scope of this work, this could in principle be tested in suitable seeding experiments. We remark that in ∼ 15 % of conformations of the single-fold double-layer N-terminal-truncated F1L2-dimer simulations, the region spanned by residues 12-21 in a chain folds back into a helix, see Figure 8a. This refolding is not observed for the full-sized F1L2 dimer. Hence, while the more rigid C-terminal region is stabilizing the fibril geometry, stacking contacts between layers involving residues 12-21 get lost without the N-terminal eleven residues, leading to an overall loss of stability and degradation of the SAA fibril. This can be seen in Figure 3c and 3d for the N-terminal truncated two-fold models F2L1- and F2L2-, which despite the additional intra-chain contacts in the C-terminal cavity (see Figure 5f) are less stable than the corresponding models F2L1 and F2L2 built from full-sized chains. The loss of stability is also seen in the MMGBSA-derived binding energies which with -7.7 (8.6) kcal/mol for F2L1- and - 8.9 (7.0) kcal/mol are much smaller in magnitude than the corresponding energies for the non-truncated F2L1 (−40.7 (11.2) kcal/mol) and F2L2 (−55.2(28.2) kcal/mol). Hence, the truncation of the N-terminal residues encourages break-up of the two protofibrils which when isolated lack the stabilizing packing contacts and easily decay, see the representative structure in Figure 8b.

**Figure 7.**
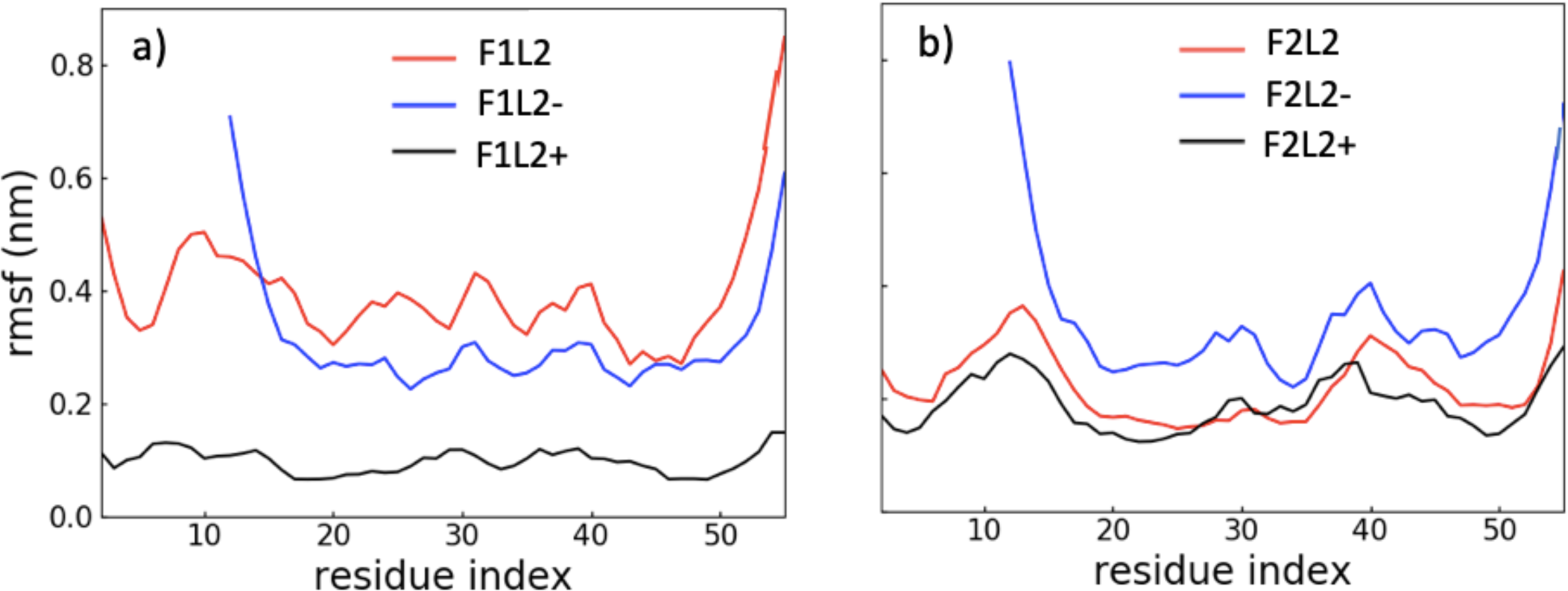
Comparison of the average RMSF of residues in full-sized and truncated (i.e. with N-terminal residues) fibril models.

**Figure 8.**
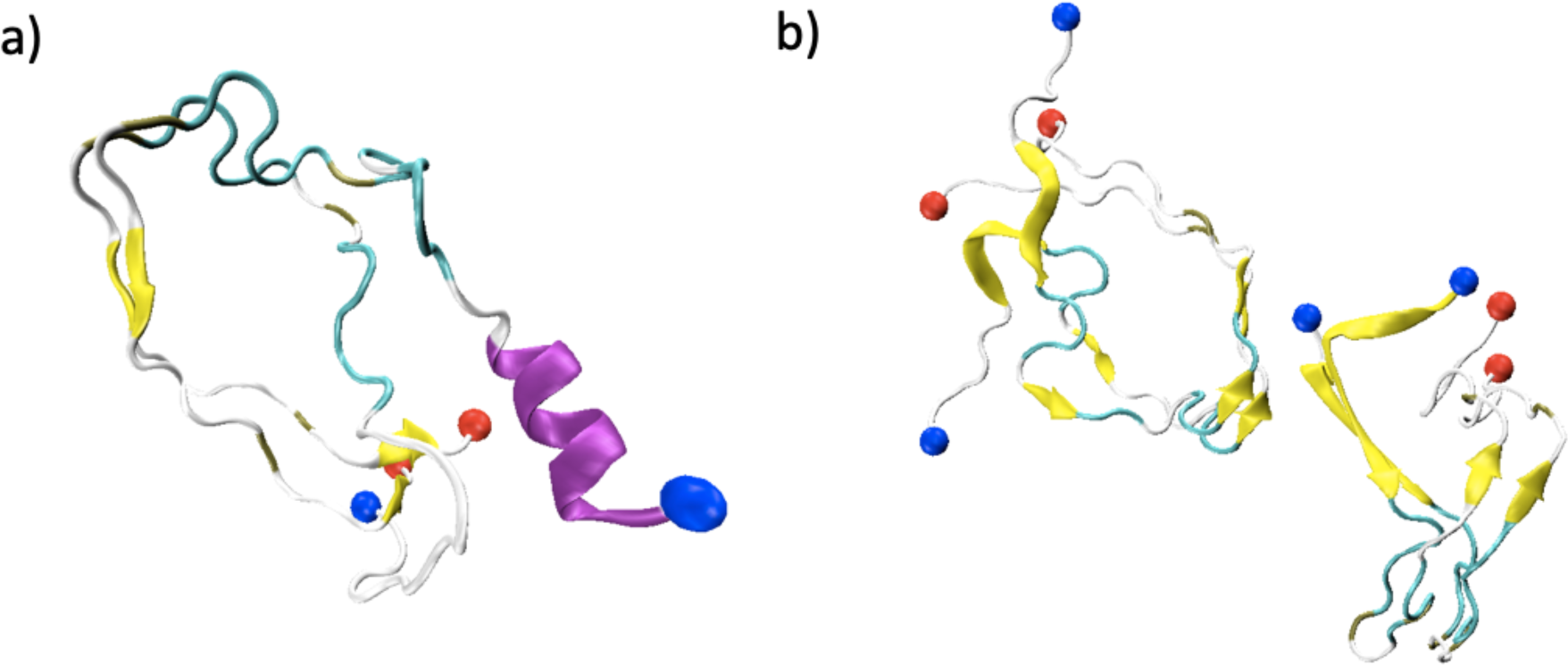
Representative structure of largest cluster seen in simulations under neutral conditions of the truncated models F1L2-(a) and F2L2-(b). Configurations from all three trajectories where clustered with a cut-off of 3 Å of systems at pH 7, and the dominant cluster contains 35% N- and C-terminus are marked by blue and red spheres, respectively. The β-strands are colored in yellow, helices in purple, turns in cyan and coil regions in gray.

The experimentally resolved SAA fibril structure contains a disordered C-terminus formed by residues 56 - 69. This region likely extends to residues 70 - 76 in the most commonly found SAA_1-76_ fibrils. Wondering about the role of this disordered region for stability of SAA fibrils, we have also performed fibril simulations where we have extended the chains in the resolved PDB-structure, assuming a disordered, random, conformation for residues 56-76. For the two-fold systems, F2L1+ and F2L2+ is the stability of the fibril region (residues 2 - 55) similar to that of the non-extended systems, see fig 2c - 2d and fig. 7b. Hence, the disordered region does not alter the stability of two-fold systems. On the other hand, for the single-fold double layer dimer F1L2+, the extension of the metastable F1L2, indicate RMSD and RMSF values in fig. 3a, 3c and 7a surprisingly a stabilization of metastable structure. This stabilization is also seen in the MMGBSA-derived binding energy approximations which with -123.5 (28.6) kcal/mol exceed the value of -59.0 (20.9) kcal/mol seen for the SAA_2-55_ fragment. This stabilization is consistent with a role of the C-terminal tail in the early stages of fibril formation but not for the stability of the fully formed fibril. We speculate that the elongated disordered C-terminal tail decreases the chances that the amyloidogenic first eleven N-terminal residues refold into the N-terminal helix, therefore stabilizing the fibril geometry at early stages.^13^

## Conclusions

SAA in its native structure is stabilized by intra-chain interactions, especially the salt-bridge between residues 1R and 9E, which stabilizes the N-terminal amyloidogenic region and prevents misfolding and aggregation. On the other hand, both packing of chains by the N-terminals and a network of ionic interactions and hydrogen bond inside the C-terminal cavity stabilize the fibril structure. We have identified as the critical size of fibrils a tetramer with two folds and two layers, which we conjecture is formed by first stacking two chains on each other, before two such dimers pack together into a two-fold structure anchored by their N-terminal regions. The critical role of the N-terminus explains the experimental observation that without the first eleven residues SAA does not aggregate into fibrils. We speculate that amyloid formation requires these residues to unravel from the N-terminal helix seen in the native structure. In earlier work^13^ we have shown that this process is aided by a disordered C-terminal region, which however, itself is not contributing to fibril stability. Once the N-terminal residues are no longer part of a helix, the now exposed segment can interact with a segment from another chain, forming a β-anchor that will be the starting point of fibril formation. While our stability analysis therefore gives already some hints on the underlying dynamics, more elaborated simulations are needed to establish in detail the mechanism of SAA fibrilization.

## Acknowledgment

The simulations in this work were done using the SCHOONER cluster of the University of Oklahoma, and XSEDE resources allocated under grant MCB160005 (National Science Foundation). We acknowledge financial support from the National Institutes of Health under research grant GM120634.

